# When touch leads vision: flexible sensory leadership during active texture exploration

**DOI:** 10.64898/2026.10.31.763287

**Authors:** Vida Arab Ahmadi, Saba Nasseri Moghaddam, Matteo Toscani, Anna Metzger

## Abstract

Vision can sample objects remotely and over a broad field, whereas touch is restricted to contacted skin and is often spatially serial however might outperform vision in resolution by transforming spatial information to a temporal code. This raises the question on how multisensory exploration is coordinated. We recorded gaze and or touch location in exploration of 60 three-dimensional surfaces with locally varying textures during visual, haptic and bimodal exploration while participants performed a same/different task, judging whether a second texture matched the first. In bimodal exploration, gaze and touch became more spatially similar than during unimodal exploration, while touch changed less than vision between unimodal and bimodal contexts. A temporal overlap analysis further showed that touch preceded gaze on average during the first 5 s of bimodal exploration. We then increased visual discriminability of the textures using cast-shadow illumination. A psychophysical pilot confirmed improved texture discrimination, and a new participant group showed a strong shift towards vision-leading exploration. Thus, sensory leadership during active visuo-haptic exploration is not fixed: touch can lead vision when texture information is important, but the balance adapts when visual information becomes more diagnostic.

## Introduction

Perception is an active process in which sensory information is acquired through movements of the sensory organs. Because we often use multiple senses to explore the same object, a central question is how these sensory systems are coordinated to gather task-relevant information. In visually guided action, the eyes typically reach the next task-relevant object before the hand, suggesting that vision plays a leading role in organising behaviour (1). This expectation is reinforced by fundamental differences in how vision and touch sample the environment. Vision can gather information at a distance and simultaneously process multiple features across a broad visual field, whereas touch requires direct contact and manual exploration is comparatively local and sequential (2–4). Moreover, the two modalities differ substantially in their anatomical capacity for spatial resolution: peak cone density in the human fovea reaches approximately 200,000 receptors/mm^2^ (5), whereas the fingertip contains approximately 241 mechanoreceptive afferents/cm^2^ (6). Consistent with this difference, at a typical reaching distance of 40 cm, normal foveal acuity corresponds to resolving spatial details on the order of 0.1 mm, compared with tactile spatial discrimination thresholds of approximately 1 mm at the fingertip (7). These differences provide a straightforward reason to expect vision to lead when the same object is explored simultaneously by sight and touch.

Yet perceptual acuity is not determined by sensory anatomy alone. Differential spatial localization (hyperacuity) increase acuity in vision (8) and touch (9). Additional acuity improvement comes from optimisation via active movement of sensory organs. In vision, finely tuned fixational eye movements enhance acuity by approximately 30%, leading to a resolution of around 82 µm at hand reach distance (10). In touch, perception of fine detail becomes even more enhanced by movement. When a finger slides across a surface, fine spatial structure is transformed into vibrations in the skin activating fast adapting tactile receptors, such as Pacinian afferents which produce a temporally patterned responses (11–13). By moving the hand across the surface, humans can distinguish surfaces with periodic wrinkles of just 760 nm wavelength and 13 nm amplitude from unpatterned surfaces (14) demonstrating sensitivity to nanoscale surface structure despite the much coarser spatial organization of cutaneous receptors and potentially outperforming vision. For judgements of surface properties, therefore, touch may provide rapidly available information that is particularly diagnostic even when vision is simultaneously available.

The question then becomes how active sampling is coordinated: where observers choose to look and where they choose to touch. In vision, the spatial allocation of gaze has long been studied in terms of saliency. Local image features contribute to making some regions more likely to attract gaze (15,16), although this stimulus-driven component interacts strongly with the observer’s task and behavioural goals (17,18). Haptic exploration has traditionally been characterised somewhat differently. Much of the literature has focused on stereotyped exploratory procedures: particular hand movements are preferentially used to judge properties such as texture, hardness, weight or temperature (19). These procedures describe how information about a given property is acquired, but not which locations within a heterogeneous surface are selected for exploration. In our previous work, we showed that this spatial selection is itself systematically related to local surface structure (20). Participants explored heterogeneous 3D-printed surfaces whose relief varied locally and independently in spatial frequency, relief height, orientation and anisotropy.

Although exploration contained a strong stimulus-independent component, reflecting stereotyped and systematic hand movements, the spatial distribution of touch duration could also be predicted from the physical relief of the particular surface being explored. Importantly, this stimulus-dependent component generalised to new participants performing a different task and even to previously unseen, naturalistic surfaces, indicating that it was not simply a consequence of a specific task strategy or stimulus set. Analysis of the model’s feature preferences suggested that participants preferentially explored edge-like, elevated and anisotropic structures, horizontal and vertical patterns, rough regions, and locations associated with rapid changes in lateral force. This provided evidence for a haptic form of saliency, in which some surface regions acquire greater exploratory priority than others. Evidence that haptic saliency also extends to 3D objects (21) suggests that such spatial prioritisation is a more general feature of active touch. These findings raise the question of how such haptic priorities are coordinated with visual selection when both senses explore the same object. When vision and touch explore the same heterogeneous surface simultaneously, do their spatial selection processes remain independent, or do they become coupled? We have recently shown that haptic exploration can alter subsequent visual exploration: after first exploring an object by touch, observers changed their fixation and saccade patterns when they later viewed the same object (22). Thus, haptic sampling can influence later visual sampling, but it remains unknown how the two exploratory systems interact when vision and touch are active at the same time.

Here we first tested whether gaze and touch became more spatially similar during bimodal than during separate unimodal exploration. We then asked whether one modality retained its unimodal exploration pattern more strongly than the other when both senses were available. Such an asymmetry would be consistent with one modality providing a spatial anchor for the other. We next asked whether this spatial coupling had a temporal direction. If one modality systematically arrived at corresponding surface regions before the other, this temporal precedence would suggest that it may be driving subsequent sampling by the other modality. We finally tested if the coordination is stable or flexibly adapts if one sense provides more information.

In a first experiment, participants explored the same heterogeneous textures used previously (20) visually, haptically and bimodally while performing a same/different texture discrimination task. We recorded gaze and index-finger position throughout exploration. Gaze and touch became more spatially similar during bimodal exploration, but touch preserved its unimodal spatial pattern more strongly than vision, suggesting that tactile sampling may provide a spatial anchor for gaze. We therefore examined the temporal relationship between the two during bimodal exploration, asking whether corresponding surface regions were sampled first by the eyes or by the finger. The relationship changed over the course of exploration: gaze tended to lead initially, but this progressively shifted towards touch-leading, with touch preceding gaze on average across the first 5 s. Together with the spatial results, this pattern suggests that touch can drive subsequent visual sampling during active texture exploration.

In a second experiment, we asked whether this apparent sensory leadership was flexible. We increased the visual discriminability of the textures using cast-shadow illumination, with the effectiveness of this manipulation to improve visual discrimination confirmed in a separate psychophysical pilot and again measured the temporal relationship between gaze and touch during simultaneous exploration. Improving the visual information substantially shifted the temporal pattern towards vision-leading exploration: gaze preceded touch through most of the first 5 s, and the transition towards touch-leading observed in the first experiment was delayed or absent. Thus, rather than reflecting a fixed dominance of either vision or touch, the direction of multisensory exploration appears to depend on the information available to each modality.

## Material and methods

### Participants

Experiment 1 included 12 adults (six women and six men; mean age = 27.5 years, range 18 – 35 years) recruited through Bournemouth University’s SONA participant system or by word of mouth. All participants were right-handed, had normal or corrected-to-normal vision, and reported no sensory or motor impairment affecting the right hand. Participants received one SONA credit or 10£ Amazon voucher per hour. All 12 participants contributed to the temporal analysis; the spatial correlation analyses included 11 participants. One participant was excluded from these analyses as eye movements were not saved in the visual only condition due to a technical issue.

Experiment 2 included eight participants (seven women and one man; mean age = 25.4 years, range 20 – 40 years), with no overlap with Experiment 1. They were all right-handed, had normal or corrected-to-normal vision, and reported no sensory or motor impairment affecting the right hand. They were recruited through Bournemouth University and compensated for their participation in the same way as for Experiment 1. A separate psychophysical pilot included other six participants and was used to establish the effect of the illumination manipulation on texture discrimination prior to the main data collection in Experiment 2.

All participants received information about the study and provided written informed consent before participation. The research was conducted in accordance with the Declaration of Helsinki and was approved by the Bournemouth University Research Ethics Committee (Ethics ID 48329).

### Stimuli

The stimulus set comprised 60 heterogeneous textured surfaces generated using the procedure described in our previous work (20). For each surface, four independent two-dimensional pink-noise maps specified the local values of relief height, spatial frequency, orientation and anisotropy across the surface (Figure 1B&C). The values in each map were mapped onto a predefined physical range for the corresponding feature. These spatially varying feature maps were then combined to generate the surface relief: a white-noise base image was filtered locally in the frequency domain using an oriented band-pass filter whose spatial frequency, orientation and anisotropy were determined by the corresponding feature values at each location, and the resulting pattern was scaled by the local height map. Thus, rather than consisting of a single homogeneous texture, each surface contained smoothly varying regions with different local texture properties. Sixty distinct surfaces were generated using different pseudorandom seeds. The resulting two-dimensional relief maps were converted into three-dimensional models by interpreting image intensity as surface depth and were then fabricated as rigid 3D-printed plates (approximately 13.97 × 13.97 cm) (20). The same set of 60 surfaces was used in the present study and, in Experiment 1, was explored under unimodal visual, unimodal haptic and bimodal visual– haptic conditions and only bimodally in Experiment 2.

**Figure 1.**
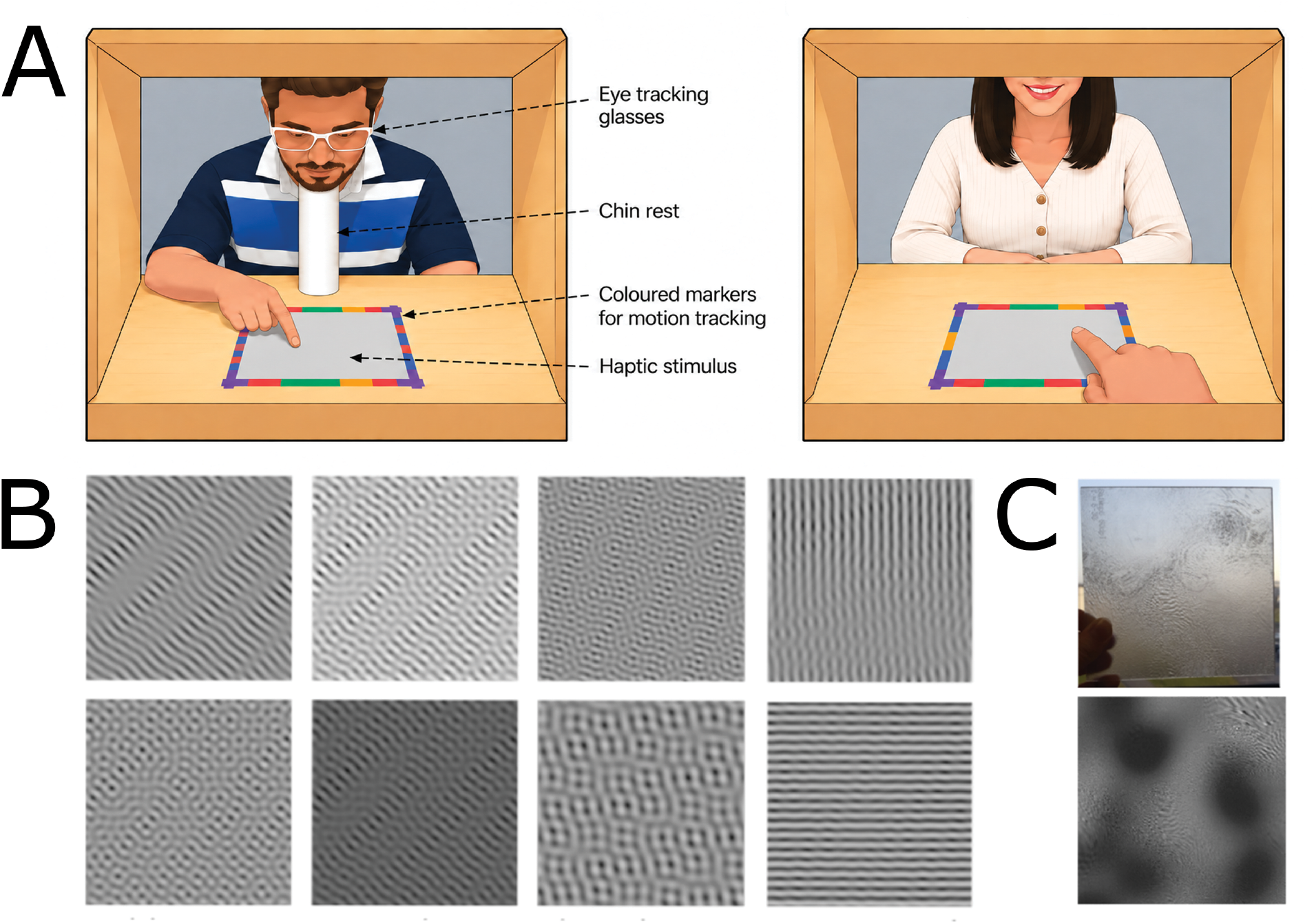
Experimental setup and stimuli. (A) Experimental setup. Left: participant seated at the apparatus wearing SMI mobile eye-tracking glasses, with the textured surface positioned horizontally in a fixed square frame at a viewing distance of approximately 40 cm. Right: schematic example image from the scene camera during bimodal exploration (B) The four features used to generate the surface reliefs: orientation, spatial frequency, height and anisotropy. Each feature is illustrated using uniform example textures, not used in the experiment, in which that feature was set to a low (top row) or high (bottom row) value while the remaining features were held constant. For the experimental stimuli, the four features varied spatially and independently across the surface according to separate pink-noise distributions. (C) Example experimental stimulus, shown as the rendered surface relief used to define the three-dimensional geometry (top) and the corresponding 3D-printed plate (bottom). Sixty different heterogeneous surfaces were generated and printed.

### Setup

The participant and the experimenter were sitting at a table on opposite sides facing each other. The stimuli were presented in a frame within wooden box (Figure 1A). The box was affixed to the table, and the frame was affixed to the box to prevent movement of the stimulus. A curtain attached to the top of the box on the participants’ side was moved down between the trials to prevent vision of the stimulus while the experimenter was placing it. To ensure constant viewing distance of 40cm participants rested the head on a chinrest. To standardise the starting position of tactile exploration, participants placed their index finger on a small finger sized support fixed to the box close to the right bottom corner of the stimulus. The frame around the stimulus was made from coloured Lego blocks to facilitate labelling of the stimulus in the video.

### Procedure

Participants performed a same/different discrimination task. The first texture served as the reference. On comparison trials, a second texture was subsequently presented and participants reported whether it was the same as or different from the reference. The experimental script randomly assigned a comparison only on 30% of trials, with equal numbers of same and different comparisons to contain the experimental session to around 1h duration. The analyses reported here concern the first, reference-texture exploration, which was available on every trial and was therefore explored in anticipation of a possible comparison. We didn’t analyse the exploration of the comparison texture as is likely conditioned by the exploration of the preceding reference texture. Exploration was not restricted.

### Eye and hand tracking

Gaze and hand movements were recorded using SensoMotoric Instruments (SMI) mobile eye-tracking glasses. The system simultaneously recorded binocular gaze and an egocentric video of the participant’s field of view through the integrated scene camera. In our recordings, gaze position was sampled at 60 Hz and the scene video was recorded at 30 frames per second, consistent with the standard SMI ETG recording configuration (23). Gaze data were exported using SMI BeGaze and expressed in the coordinate system of the scene video.

Fingertip and stimulus position was extracted from the scene videos using DeepLabCut, a markerless pose-estimation method based on supervised deep learning (24). We followed the standard DeepLabCut workflow (25). For each participant and sensory condition, training frames were sampled from two randomly selected exploration videos; 10 frames were selected and manually labelled with the position of the index fingertip and the four corners of the stimulus.

These labelled frames were used to train the network, which was subsequently applied to the complete set of scene-video frames to estimate the five landmark positions automatically. Tracking confidence was provided by DeepLabCut for each estimate, and fingertip estimates with likelihood below 0.9 were excluded from analysis.

Because gaze position and the DeepLabCut landmarks were defined in the same scene-camera image, the four tracked stimulus corners provided a direct mapping between image coordinates and coordinates on the physical texture. For each recording, the corners were used to compute a projective transformation from the scene image onto the two-dimensional stimulus plane. The same transformation was then applied to gaze and index-fingertip positions, placing both measures in a common surface-centred coordinate system and allowing their spatial distributions to be compared directly.

### Spatial exploration maps

Analysis followed previous work on haptic saliency using these stimuli (20). For each participant, stimulus and condition, gaze and touch samples were accumulated in a 330 × 330 surface map. Repeated samples at the same location were accumulated so the maps represented spatial sample density. Maps were smoothed with a Gaussian kernel (sigma = 16.5 pixels) and normalised to sum to one. Spatial similarity was quantified with Pearson correlation across corresponding pixels. Two participant-level comparisons were defined a priori by the scientific questions. First, we compared gaze-touch similarity during separate unimodal explorations with gaze-touch similarity during bimodal exploration. Second, we quantified the stability of each modality across contexts by correlating unimodal touch with bimodal touch, and unimodal vision with bimodal vision. Differences between the two correlations in each question were tested with two-sided paired t-tests across participants.

### Temporal overlap analysis

Temporal analyses used bimodal trials. Gaze and finger trajectories were expressed in the scene-video time base. The texture surface was divided into a 10 x 10 spatial grid. For each trial we constructed a gaze-time x touch-time matrix whose entries indicated whether gaze and touch occupied the same spatial cell at the corresponding pair of times. Each non-empty trial matrix was divided by its total overlap before participant-level aggregation so that trials contributed equally. For each gaze time point, we identified the touch time point (or the mean of tied time points) with the maximal aggregate spatial overlap. Temporal offset was defined as touch time minus gaze time: negative values therefore indicate that touch occupied the corresponding region before gaze, whereas positive values indicate that gaze came first. For the experiment comparison, both gaze and touch were restricted to the first 5.0 s before the overlap matrix was computed. Each participant’s mean offset across the full 5-s common window was first tested against zero using a two-sided one-sample t-test. To examine how the temporal relationship evolved during exploration, offsets were additionally averaged within five consecutive 1-s bins (0–1, 1–2, 2–3, 3–4 and 4–5 s). Mean offsets in each bin were tested against zero using two-sided one-sample t-tests, with Holm correction across the five tests within each experiment. The time-binned data were further analysed with a mixed 2 x 5 ANOVA, with Experiment (Experiment 1 vs Experiment 2) as a between-subject factor and Time as a within-subject factor. Experiment 1 and Experiment 2 were also compared separately within each 1-s bin using two-sided Welch independent-samples t-tests, with Holm correction across the five comparisons.

### Experiment 2: improved visual discriminability

Experiment 2 repeated bimodal texture exploration in a new group of eight participants under an illumination configuration designed to make surface relief more visually discriminable. The manipulation used pronounced cast shadows rather than the ambient-light condition used in the visibility pilot. The stimuli were illuminated with a directional lamp positioned laterally at an angle of approximately 45° to the surface, producing pronounced cast shadows across the textured relief. Oblique illumination is known to enhance image variations generated by three-dimensional surface structure, including shading gradients and cast shadows, which provide strong cues to surface roughness and relief (26,27).

The effectiveness of the cast-shadow manipulation was assessed in a separate psychophysical pilot using the same visual same/different discrimination task as in the visual condition of Experiment 1. Participants viewed a reference texture and judged whether the second comparison texture was the same as or different from the sample. Each participant completed 30 trials under ambient illumination and 30 trials under cast-shadow illumination, with each texture presented for 5 s. Discrimination accuracy increased from 73.9% under ambient illumination to 84.4% under cast-shadow illumination, a mean improvement of 10.6 percentage points. This improvement was significant in a two-sided paired-samples *t*-test, *t*(5) = 3.63, *p* = .015, confirming that the cast-shadow illumination increased the visual discriminability of the textured surfaces.

### Experiment comparison and time-resolved statistics

Because Experiment 1 and Experiment 2 contained different participants, between-experiment tests treated Experiment as a between-subject factor. The primary overall comparison used a two-sided *t*-test on participant mean offsets over the common 5 s interval.

For the time-resolved analysis, the 5 s interval was divided into five non-overlapping 1 s bins. A 2 (Experiment: Experiment 1, Experiment 2) x 5 (Time) mixed-design ANOVA tested the main effects of Experiment and Time and their interaction. Time was the repeated factor and the standard univariate test assumed sphericity. Post hoc *t*-tests compared experiments within each time bin, and one-sample *t*-tests compared each bin with zero within each experiment. Holm correction controlled family-wise error across the five bins within each family of comparisons.

## Results

### Experiment 1: Bimodal exploration increases spatial coupling

Visual and haptic exploration appeared spatially distinct in the unimodal conditions, but became noticeably more similar during simultaneous visuo-haptic exploration (Examples in Figure 2A). Gaze and touch were only weakly correlated when the visual and haptic explorations were performed separately (mean *r* = 0.012). When both modalities were available simultaneously, their spatial exploration maps were substantially more similar (mean *r* = 0.241; Figure 2B). The participant-level increase was 0.229 correlation units (95% CI 0.115 to 0.343), *t*(10) = 4.46, *p* = 0.001.

**Figure 2.**
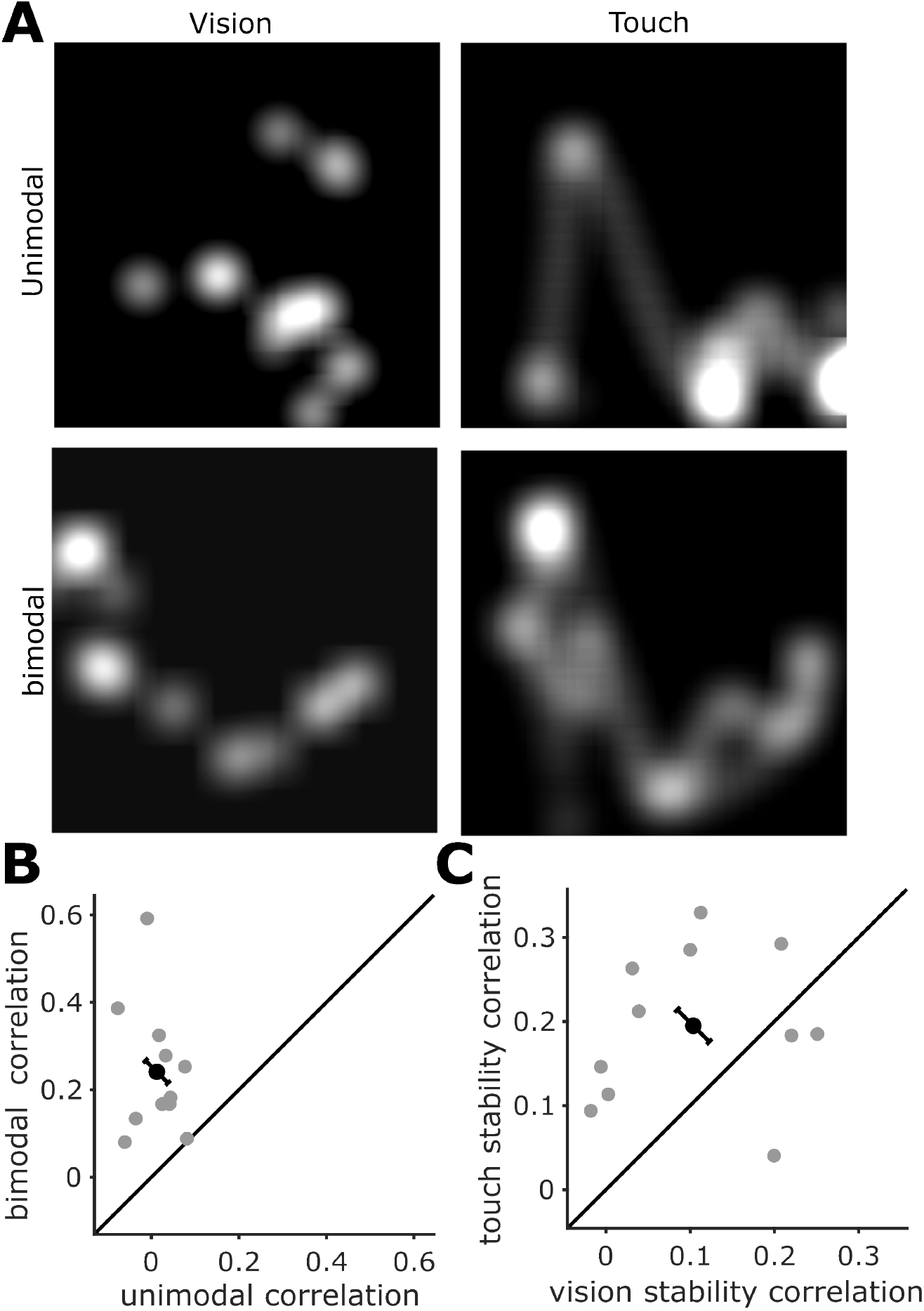
Spatial coupling of gaze and touch during texture exploration. (A) Example gaze- and touch-density maps from one participant. Columns show visual exploration (left) and haptic exploration (right); rows show unimodal exploration (top) and bimodal exploration (bottom). Brighter regions indicate locations sampled more frequently. (B) Participant-level Pearson correlations between gaze and touch density maps during unimodal exploration (x-axis) and bimodal exploration (y-axis). (C) Stability of modality-specific exploration patterns across sensory contexts: the x-axis shows the correlation between unimodal and bimodal gaze maps, and the y-axis the corresponding correlation between unimodal and bimodal touch maps. In B and C, grey points represent individual participants, the black point represents the group mean, and the black error bar shows ±1 SEM of the paired difference between the two quantities, plotted perpendicular to the unity line. The diagonal line (unity line) indicates equality between the x- and y-axis measures; points above the line indicate a higher bimodal than unimodal gaze–touch correlation in B, and greater touch than vision stability in C.

### Touch is more stable than vision across sensory contexts

We next asked whether one modality retained a more similar spatial pattern when moving from unimodal to bimodal exploration. Touch maps were more stable across contexts (mean *r* = 0.195) than gaze maps (mean *r* = 0.104; Figure 2C). The mean difference was 0.091 (95% CI 0.006 to 0.176), *t*(10) = 2.40, *p* = 0.038. Thus, simultaneous exploration increased cross-modal coupling while the haptic exploration pattern changed less than the visual pattern. This asymmetry is consistent with touch providing a spatial constraint on gaze.

### Touch precedes gaze during the later phase of texture exploration

The temporal structure is illustrated by the overlap matrix for an example participant (Figure 3A). Early in exploration, the strongest spatial overlap in time lay close to simultaneity (the diagonal) and extended towards later finger times (maximum overlap below the diagonal), consistent with gaze initially preceding touch and touch dominating in the remaining exploration i.e. the finger increasingly reaching corresponding surface locations before the eyes did. Across the common first 5 s of Experiment 1, the mean touch-minus-gaze offset was -0.297 s (95% CI -0.578 to - 0.016), *t*(11) = -2.32, *p* = 0.04, indicating that corresponding touched locations tended to be reached before they were fixated (Figure 3B). The time-resolved pattern was not uniform: gaze led early (as indicated by positive touch – gaze values), whereas touch led later in the exploration (as indicated by negative touch – gaze values).

**Figure 3.**
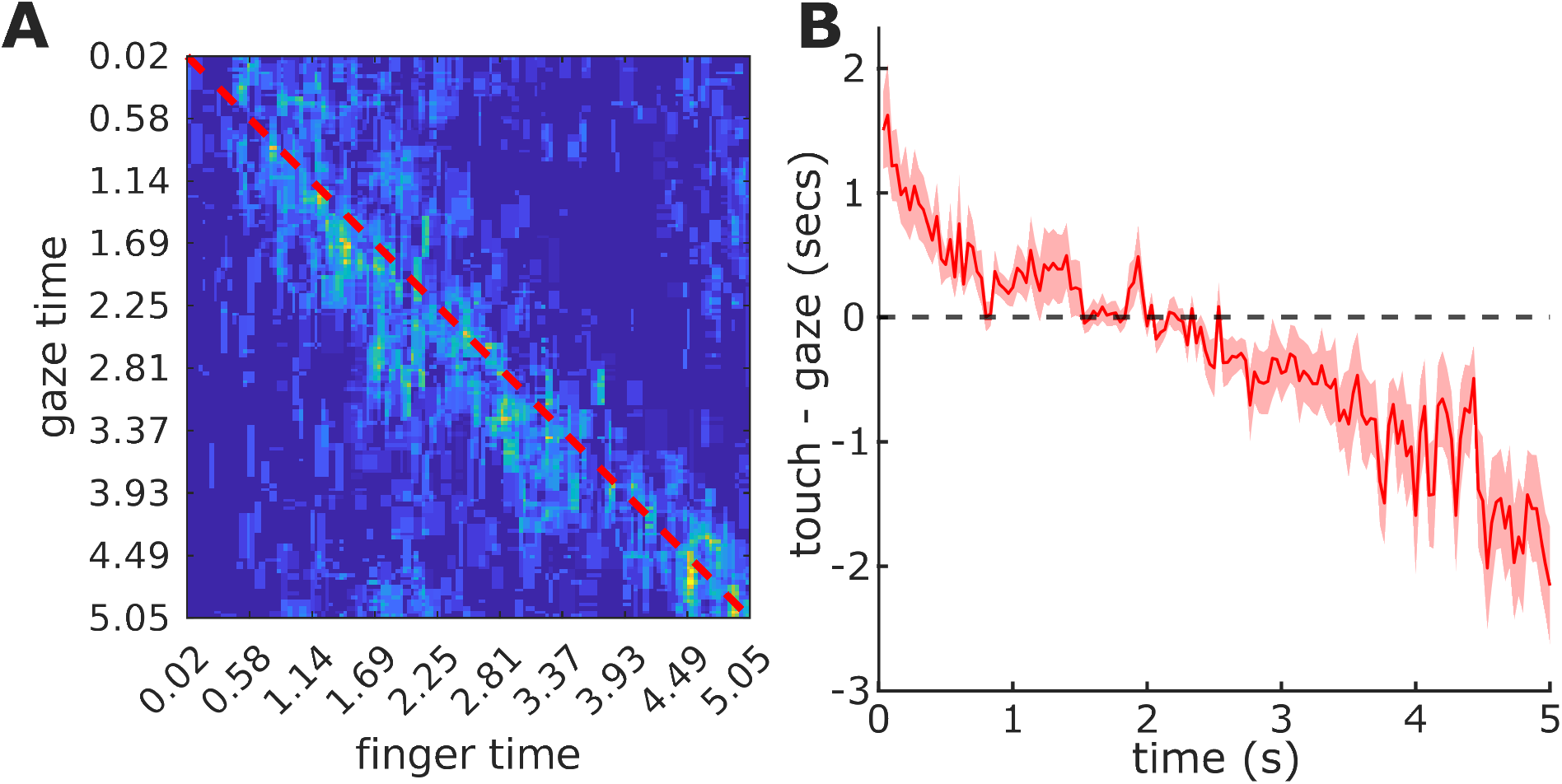
Temporal relationship between gaze and touch during bimodal exploration in Experiment 1. (A) Gaze-time × finger-time overlap matrix for an example participant, averaged across trials. Colour indicates the relative spatial overlap between gaze and touch for each pair of time points; the dashed diagonal marks simultaneous sampling. (B) Mean temporal offset between touch and gaze across participants over the first 5 s of exploration. The offset is defined as touch time minus gaze time; positive values indicate that gaze preceded touch, whereas negative values indicate that touch preceded gaze. The shaded region shows ±1 SEM.

After Holm correction across the five 1 s bins, Experiment 1 showed a positive offset during 0-1 s (mean = 0.650 s, adjusted p < 0.001), no reliable offset during 1-2 s or 2-3 s, and reliable touch-leading offsets during 3-4 s (mean = -0.742 s, adjusted *p* = 0.019) and 4-5 s (mean = -1.362 s, adjusted *p* = 0.004). This progression suggests that touch did not simply precede vision from trial onset; rather, tactile exploration became temporally leading as the texture was explored.

### Experiment 2: Improved visual discriminability shifts sensory leadership towards vision

Under the improved-visibility condition, the temporal relationship shifted strongly towards vision. Across the same first 5 s used for analysis in Experiment 1, Experiment 2 showed a positive mean touch-minus-gaze offset of 1.394 s (95% CI 1.227 to 1.562), *t*(7) = 19.64, *p* < 0.001. The between-experiment shift was 1.691 s (95% CI 1.382 to 2.001), Welch *t*(16.39) = 11.57, *p* < 0.001.

The time-resolved mixed ANOVA confirmed that the experiments differed not only in their average offset but also in how the offset developed over time (Figure 4). There were effects of Experiment, *F*(1,18) = 103.62, *p* < 0.001, partial eta-squared = 0.852, and Time, *F*(4,72) = 98.65, *p* < 0.001, partial eta-squared = 0.846, together with an Experiment x Time interaction, *F*(4,72) = 11.21, *p* < 0.001, partial eta-squared = 0.384.

**Figure 4.**
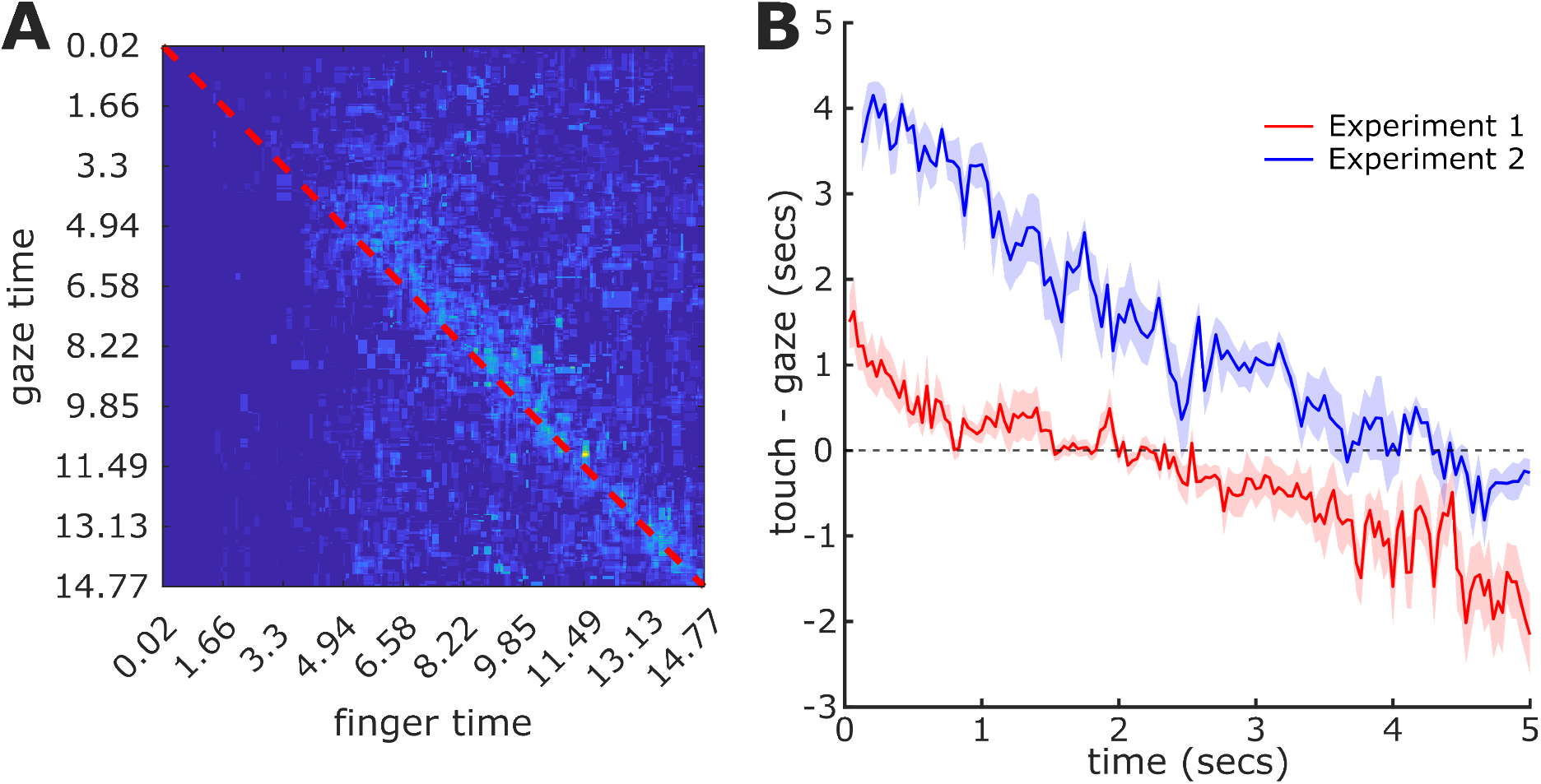
Temporal relationship between gaze and touch under improved visual discriminability. (A) Gaze-time × finger-time overlap matrix for Experiment 2, example participant. Colour indicates the relative spatial overlap between gaze and touch for each pair of time points; the dashed diagonal marks simultaneous sampling. (B) Mean touch-minus-gaze offset during the first 5 s of exploration for Experiment 1 (red) and Experiment 2 (blue). Positive values indicate that gaze preceded touch, whereas negative values indicate that touch preceded gaze. Shaded regions show ±1 SEM across participants; the horizontal dashed line indicates zero offset.

Holm-corrected post hoc tests showed a larger (more vision-leading) offset in Experiment 2 than Experiment 1 in every 1 s bin (all adjusted *p* <= 0.0024). Within Experiment 2, gaze led reliably during each of the first four bins (0-4 s; all adjusted *p* <= 0.048), whereas the final 4-5 s bin did not differ from zero (mean = -0.162 s, adjusted *p* = 0.122). Thus, the touch-leading phase evident late in Experiment 1 did not emerge reliably when visual surface information was enhanced.

## Discussion

The central result of this study is that there is no fixed vision-first hierarchy during active visual-haptic texture exploration. Under the original laboratory illumination, visual and haptic exploration became more spatially coupled when both modalities were available, the touch pattern was more stable than the gaze pattern across unimodal and bimodal contexts, and touch preceded gaze during most of the exploration except for the initial phase, indicating that tactile sampling became temporally leading. When visual surface structure was made more discriminable by cast-shadow illumination, the temporal relationship shifted strongly towards vision. Together, these results suggest flexible sensory leadership: touch can organise where and when vision samples a textured surface, but its leading role is reduced when visual information becomes more diagnostic.

A touch-led contribution is initially counterintuitive. Vision has access to a much larger region of space, does not require physical contact and is efficient at extracting multiple features in parallel (2). In everyday object-directed actions, gaze typically precedes manual contact (1). Haptic exploration, in contrast, is constrained by the size and location of the contacting skin and often requires sequential movements across the object (3,4). If sensory leadership reflected sampling range alone, vision should therefore dominate. Texture provides a plausible mechanism for reversing this expectation. Sliding contact converts fine spatial surface structure into high-frequency skin vibrations, and the tactile system preserves behaviourally relevant information in precise temporal response patterns (11–13). The present task required participants to acquire information that could later support a same/different judgement of textured surfaces. Under these circumstances, touch may rapidly detect diagnostic surface structure, while vision subsequently samples those regions to establish their local spatial arrangement or to integrate them with broader visual information as precise localisation via touch might be less accessible when involving the temporal code for resolving fine detail. This interpretation is also consistent with classic evidence that the relative contribution of vision and touch depends on the judged texture dimension rather than following a fixed dominance hierarchy (28).

The spatial results should nevertheless be interpreted carefully. Greater gaze-touch correlation in the bimodal condition demonstrates coupling, not direction. Similarly, the greater stability of the touch map is consistent with, but does not prove, the idea that haptic exploration constrains gaze. The temporal analysis provides stronger directional information because it asks whether spatially corresponding touch events tend to occur before or after gaze.

The visibility manipulation provides an important test of flexibility. Multisensory cue weighting is known to depend on relative sensory reliability (29), and visual-haptic texture judgements can reverse their apparent modality dominance depending on the information required (28). Here, a simple change in illumination improved visual texture discrimination and substantially changed the exploration dynamics. The result argues against a hard-wired touch-first strategy for textures. Instead, the system appears to adapt the ordering of active sampling to the information available from each modality.

The present experiments focused primarily on texture, a property for which haptic information can be particularly diagnostic. Objects, however, also have geometric properties such as shape, for which vision is often the more effective modality (3,30). Spatially selective haptic exploration is not restricted to texture. In recent work with three-dimensional objects, it was found that the locations selected for touch could be predicted from local curvature features, indicating that particular geometric regions have greater haptic salience during shape exploration (21). Moreover, differences in touching strategy were associated with performance in an object-recognition task, suggesting that where an object is touched is behaviourally relevant for extracting shape information. These findings make shape a particularly interesting test case for sensory leadership. Our previous study further suggests that tactile leadership may not be restricted to texture. In that study, participants first explored three-dimensional object shapes haptically and subsequently viewed the same objects; prior touch altered later fixation and saccade patterns, producing a broader and more spatially distributed pattern of visual exploration that resembled the exploration expected during touch (22). This raises the possibility that haptic exploration can also lead subsequent visual sampling when shape, rather than texture, is the relevant object property. Because vision and touch were sequential in that study, however, temporal leadership could not be tested directly. Whether touch can also become temporally leading during simultaneous exploration of shape therefore remains an open question. More generally, natural objects normally provide shape and texture together rather than in isolation. An important next step will be to determine how visual and haptic leadership is negotiated when both kinds of information are simultaneously available, and whether the leading modality changes dynamically according to which object property is most informative at a given moment.

More broadly, the present findings highlight a distinction between integrating sensory information and coordinating its acquisition. Much of the visual–haptic literature has asked how estimates obtained through vision and touch are combined, for example whether each modality is weighted according to its reliability or whether a common multisensory representation is formed (29,31). Such approaches address what happens once sensory evidence is available. Active perception poses an additional problem: the observer must determine where and when that evidence will be acquired. Both vision and touch are active senses in this respect. Eye movements determine which parts of an object receive high-resolution visual processing, while hand movements determine which parts of an object generate tactile input. Active-sensing theories therefore treat perception and sampling action as a closed loop in which current information determines the next sensory action (32).

These two aspects of the problem have largely been studied separately. Research on eye–hand coordination has characterized in considerable detail how gaze and the hand are coordinated during reaching and object manipulation, with gaze typically arriving at task-relevant locations before the hand (1). In that literature, however, the hand is primarily an effector whose movement is visually guided. Conversely, research on active touch has shown that hand movements are specifically organized to acquire particular object properties and that the spatial distribution of touch can itself depend on local surface structure (4,19,20). Here the hand has the role of a sensor, but the simultaneous allocation of visual sampling has generally not been measured. The present study directly examines how gaze and fingertip sampling are coordinated when vision and touch explore the same object simultaneously.

An especially relevant precursor is the study by Liesker, Brenner and Smeets (33), who asked participants to search visually and haptically at the same time. Combined search was faster than search with either modality alone, but less efficient than the optimal strategy in which vision and touch would divide the search space and inspect different locations. Their results were consistent with the two modalities operating in parallel while partly sampling overlapping locations. The present findings suggest that such overlap need not simply reflect redundant search. At a finer spatial and temporal scale, one modality may sample a location first and the other subsequently visit the same region. Thus, what appears as suboptimal overlap when measured only through overall search time may in part reflect coordinated multisensory sampling. Continuous tracking of gaze and hand trajectories is required to distinguish these possibilities. Our results provide evidence for such coordination: rather than behaving as two wholly independent searches, gaze and touch became more spatially coupled when they sampled the same textured surface, and that coupling acquired a changing temporal direction over the course of exploration.

Crossmodal-attention studies provide another important link. Haptic information can bias subsequent visual selection even when the tactile and visual targets are physically separate. For example, holding an unseen shape accelerated eye movements towards visually presented objects with a corresponding shape (34), demonstrating haptic guidance of overt visual attention. Similarly, prior haptic exploration of object shape altered the distribution of subsequent visual exploration (22). These studies establish that information acquired by touch can change where the visual system allocates attention, but they do not reveal how the two sensors coordinate their sampling when they are simultaneously exploring the same physical surface. The present results extend this literature from crossmodal influences on later or spatially separate visual selection to the continuous coordination of gaze and touch during ongoing exploration.

To our knowledge, this is the first direct characterization of the spatial and temporal coupling of visual and haptic sampling trajectories during simultaneous exploration of the same heterogeneous surface. Addressing this question requires more than measuring perceptual judgements or total exploration times: gaze and fingertip positions must be recovered continuously and expressed in a common object-centred coordinate system. This has become considerably more practical with recent developments in mobile eye tracking and markerless, deep-learning-based motion tracking, which allow hand position and object geometry to be recovered from naturalistic video (24).

Distinguishing sensory integration from sensory sampling also changes how the illumination result should be interpreted. Improved visual information did not simply increase the weight given to vision after visual and haptic information had been acquired. It changed how information was acquired in the first place: under cast-shadow illumination, gaze remained ahead of touch for longer, indicating that vision more often selected surface locations before they were sampled haptically. In other words, the manipulation altered the temporal ordering of exploration - which modality sampled first and which tended to follow - rather than merely changing the relative contribution of two already-available sensory estimates. Sensory reliability may therefore influence multisensory perception at an earlier stage than usually considered in cue-integration accounts, by determining which modality selects information first and which modality subsequently samples corresponding regions.

The broader implication is that multisensory exploration may be organized as a coupled active-sensing system. Which sense leads need not be specified in advance. Vision can provide the first survey of an object, touch can subsequently identify locally informative structure and recruit visual sampling to corresponding locations, and the balance can shift again when the quality of visual information changes. Together with the evidence that prior haptic exploration can reorganize later visual sampling of shape, the present results suggest that such crossmodal control may extend beyond texture. Determining how this coordination operates when natural objects simultaneously provide texture, shape and other material properties will be important for understanding multisensory perception under the conditions in which it normally occurs.

In conclusion, our results show that visual–haptic exploration is not governed by a fixed sensory hierarchy. Instead, gaze and touch coordinate their sampling dynamically, with the modality that leads depending on the information available and changing over the course of exploration. This extends multisensory research beyond the integration of already acquired signals to the control of information acquisition itself. By jointly tracking gaze and fingertip trajectories on the same surface, we show that multisensory perception involves not only combining sensory evidence, but also coordinating which sense samples first, where it samples, and which sense follows

## Data, code and declarations

The analysis code and data underlying the reported results will be deposited in GitHub upon acceptance of the manuscript.

The research was supported by the Royal Society RG\R1\241159 – Research Grant. We thank Knut Drewing for helpful discussions. The authors declare no competing interests.

